# EHF is essential for epidermal and colonic epithelial homeostasis and suppresses *Apc*-initiated colonic tumorigenesis

**DOI:** 10.1101/2021.03.01.433470

**Authors:** Camilla M. Reehorst, Rebecca Nightingale, Ian Y. Luk, Laura Jenkins, Frank Koentgen, David S. Williams, Charbel Darido, Fiona Tan, Holly Anderton, Michael Chopin, Kael Schoffer, Moritz F. Eissmann, Michael Buchert, Dmitri Mouradov, Oliver M Sieber, Matthias Ernst, Amardeep S. Dhillon, John M. Mariadason

## Abstract

**Background:** Ets homologous factor (EHF) is a member of the epithelial-specific Ets (ESE) transcription factors. EHF is specifically expressed in epithelial tissues, however its role in development and epithelial homeostasis is largely uncharacterized.

**Methods:** We generated a novel mouse strain in which the Ets DNA binding domain (exon 8) of Ehf was flanked by loxP sites (*Ehf*^*Lox/Lox*^). To inactivate Ehf in the whole body, *Ehf*^*Lox/Lox*^ mice were crossed to *CMV^Cre^* mice, which were then bred out to generate germline *Ehf* null (*Ehf*^*−/−*^) mice. To inactivate Ehf specifically in the intestinal epithelium, *Ehf^Lox/Lox^* mice were bred to tamoxifen-inducible *Villin^Cre-ERT2^* mice. *Ehf^Lox/Lox^* mice were also crossed to tamoxifen-inducible *Cdx2^CreERT2^*; *Apc*^*Lox/+*^ mice to determine the impact of Ehf deletion on Apc-initiated colon cancer development.

**Results:** Transcripts encoding the Ets binding domain of EHF were effectively deleted in all tissues in *Ehf*^*−/−*^ mice. *Ehf*^*−/−*^ mice were born at the expected Mendelian ratio, but showed reduced body weight gain and developed a series of pathologies during their lifespan that led the majority of *Ehf*^*−/−*^ mice to reach an ethical endpoint within one year of age. Most prominent of these were the development of papillomas in the chin, and abscesses in the preputial glands (males) or vulvae (females) which showed evidence of Staphylococcus and Proteus infection. Consistent with the development of papillomas, the epidermis of *Ehf*^*−/−*^ mice showed evidence of mild hyperplasia. A subset of *Ehf*^*−/−*^ mice also developed cataracts and corneal ulcers. EHF is highly expressed in the colonic epithelium and *Ehf*^*−/−*^ mice displayed increased susceptibility to dextran sodium sulphate-induced colitis. This phenotype was confirmed in intestinal-specific Ehf knockout mice, and histopathological analyses revealed reduced numbers of goblet cells and extensive transcriptional reprogramming in the colonic epithelium. Finally, colon-specific deletion of *Ehf* enhanced *Apc*-initiated adenoma development, unveiling a novel, tumour suppressive role for EHF in colorectal cancer.

**Conclusion:** The Ets DNA-binding domain of EHF is essential for post-natal homeostasis of the epidermis and colonic epithelium, and functions as a tumour suppressor in the colon.

## Introduction

Ets homologous factor (*EHF*) is a member of the Ets family of transcription factors, which consists of 35 members in humans. All Ets family members share a conserved winged helix-turn-helix *ETS* DNA binding domain, through which they bind to 5’-GGAA/T-3’ sequence elements in DNA (Brembeck et al., 2000). Among these, EHF is a member of the ESE subfamily, comprising ELF3, ELF5, EHF and SPDEF, which are grouped together based on their common epithelial-specific expression profile. All ESE members also share an N-terminal Pointed (PNT) domain comprising ~80 amino acids (aa) (Seidel and Graves, 2002), which is involved in protein-protein interactions, kinase docking, RNA-binding and lipid molecule interactions, and can have strong transactivation activity (Piggin et al., 2016).

*EHF* is located on chromosome 11p13 and encodes a 300 amino acid protein. Expression of EHF in normal human tissues is highest in the salivary gland, oesophagus, vagina, prostate, colon, skin, bladder and breast (Luk et al., 2018). The role of EHF in tumour progression has been relatively well studied, with loss of expression and a potential tumour suppressive role suggested in prostate (Albino et al., 2012), pancreatic (Zhao et al., 2017) and oesophageal squamous cell carcinoma (Wang et al., 2015), where EHF loss has been shown to promote epithelial-to-mesenchymal transition and cell migration. Conversely, EHF has been shown to promote proliferation in cell line models of gastric (Shi et al., 2016), thyroid (Lv et al., 2016) and ovarian cancer (Cheng et al., 2016).

In contrast to its role in cancer, few studies have investigated the role of EHF in normal tissue homeostasis. In line with its high expression in the skin, a study using primary cell cultures suggested a key role for EHF in keratinocyte differentiation (Rubin et al., 2017). Profiling of enhancer regions in keratinocytes identified the *GGAA* Ets binding motif as the most enriched transcription factor binding site, and EHF knockdown in organotypic human epidermal tissue revealed that EHF regulates ~400 genes, including several associated with keratinocyte differentiation (Rubin et al., 2017). EHF is also highly expressed in the intestinal epithelium, and a role for EHF in maintaining the colonic stem cell compartment was recently proposed (Zhu et al., 2018). However, the impact of *Ehf* deletion on homeostasis in these tissues has not been systematically addressed *invivo*.

In this study we induced whole body deletion of the Ets DNA binding domain of EHF. *Ehf* knockout (*Ehf*^*−/−*^) mice were viable and fertile but developed a range of pathologies over their lifespan including infections in the preputial glands of males or vulvae of females, skin papillomas, and less frequently corneal ulcers. *Ehf* deletion also impacted intestinal epithelial homeostasis, and increased susceptibility to DSS-induced acute colitis. Finally, colon-specific deletion of *Ehf* increased Apc-initiated adenoma development, revealing a previously unknown tumour suppressive function of EHF in the colon. These findings identify a key role for EHF in maintaining epidermal and intestinal epithelial homeostasis, and suppression of tumour formation in the colon.

## Methods

### Mice

*Ehf*^*Lox/Lox*^ mice were generated by flanking exon 8 and the coding region of exon 9 with *loxP* sites in C57Bl/6 embryonic stem (ES) cells. The targeting vector contained a neomycin cassette for selection in ES cells and was flanked by *FRT* sites for Flp-recombinase-mediated removal of the neomycin cassette. ES cell clones were selected based on neomycin resistance, screened by Southern hybridisation, and correctly targeted clones were microinjected into C57BL/6 murine blastocysts, and implanted into pseudo-pregnant females. Resulting chimeras were bred to wildtype C57Bl/6 mice to generate mice heterozygous for the floxed allele. Heterozygous mice were then crossed with OzFLP mice to induce removal of the neomycin cassette. Mice heterozygous for the floxed *Ehf* allele were subsequently mated to generate homozygous *Ehf*^*Lox/Lox*^ mice. *Ehf*^*Lox/Lox*^ mice were subsequently mated with C57Bl/6 Tg(CMV-cre) mice (Original Source: Mouse Genetics Cologne (MGC) Foundation, provided by Warren Alexander, Walter and Eliza Hall Institute, Melbourne, Australia) to induce constitutive *Ehf* deletion. *CMV-cre*^*e*^ was then bred out of the colony, and germline *Ehf*^*−/−*^ and *Ehf^+/+^* control progeny obtained by *Ehf^+/−^* heterozygous mating.

To selectively induce *Ehf* deletion in intestinal epithelial cells, *Ehf*^*Lox/Lox*^ mice were crossed to C57Bl/6 *Villin*^*CreERT2*^ mice (provided by Dr. Sylvie Robine)(el Marjou et al., 2004)) to generate *Ehf*^*Lox/Lox*^;*Villin^CreERT2^* mice. At 4-5 weeks of age, mice were intraperitoneally injected with a total of 9 mg of tamoxifen solution (10mg/mL) dissolved in sunflower seed oil, over four consecutive days. Control mice received sunflower seed oil only. To determine the impact of *Ehf* deletion on Apc-initiated colon tumour development, *Ehf*^*Lox/Lox*^ mice were bred to *Cdx2*^*CreERT2*^;*Apc*^*Lox/+*^ mice to generate *Ehf*^*Lox/Lox*^; *Apc*^Lox/+^;*Cdx2*^*CreERT2*^ (*Ehf*^*CKO*^ *Apc*^*CKO/+*^) and *Ehf^+/+^*; *Apc*^*Lox/+*^; *Cdx2*^*CreERT2*^(*Ehf*^*WT*^ *Apc*^*CKO/+*^) control mice. Apc^Lox/Lox^ (580S) (Shibata et al., 1997) and Cdx2-CreER mice (Hinoi et al., 2007) have been described previously. At 4-5 weeks of age, mice were treated with 9 mg of tamoxifen as above. All mice were maintained on pure C57Bl/6 backgrounds. All animal breeding and procedures were performed in accordance with approval obtained from the Animal Ethics Committee, Austin Health (Melbourne). All animals were housed in open-top boxes with basic enrichment, and provided with standard mouse chow and drinking water *ad libitum*. Primers used for DNA genotyping were as follows: Intron 7 (outside floxed region)-3’UTR exon 8 (outside floxed region) *Ehf* (F): GTCCAAAATGAAGCCCAGGGTA, *Ehf* (R): 5’-CGTCCGGTTCTTCATTGATCAG. Intron 7 (inside floxed region)-3’UTR exon 8 (outside floxed region) *Ehf* (F): 5’-TGTGTCTTGCTTTCCACCAG, (R): 5’-CGTCCGGTTCTTCATTGATCAG. *CMV^Cre^* (F): 5’-CTGACCGTACACCAAAATTGCCTG, (R): 5’-GATAATCGCGAACATCTTCAGGTT. *Villin^CreERT2^* (F): 5’-CAAGCCTGGCTCGACGGCC, (R) 5’-CGCGAACATCTTCAGGTTCT. *Apc* (F): 5’-GTTCTGTATCATGGAAAGATAGGTGGTC, (R): 5’-CACTCAAAACGCTTTTGAGGGTTGATTC. *Cdx2^CreERT2^* (F): 5’-CATGGTGAGGTCTGCTGATG, (R): 5’-CATGTCCATCAGGTTCTTGC.

### Periodic Acid-Schiff and Alcian Blue stains

For detection of goblet cells, deparaffinised sections were stained with Alcian blue 8GX (ProSciTech, Australia) for 15 minutes. Slides were then rinsed thoroughly in water and stained in 0.5% periodic acid in distilled water (Sigma-Aldrich, USA) for 5 minutes. Slides were then washed and stained with Schiff’s reagent (VWR International, USA) for 10 minutes, counterstained with haematoxylin, dehydrated through serial ethanol and xylene washes, and mounted using DPX (Sigma Aldrich, USA).

### Immunohistochemistry

Deparaffinised and rehydrated formalin fixed paraffin-embedded (FFPE) sections were treated with 3% H_2_O_2_ (Science Supply Associates, Australia) for 10 minutes at room temperature and Antigen retrieval performed by incubation in citrate buffer (10mM Sodium citrate, 0.05% Tween, pH 6.0) at 100°C for 30 minutes. Slides were then washed in Tris-buffered saline with tween (TBST, 0.05M Tris, 0.9% NaCl, 0.05% Tween, pH 7.6), blocked and stained with primary antibody overnight at 4°C. Antibodies used in IHC analyses were: Anti-Chromogranin A (ab85554, Abcam USA, 1:500), Lysozyme/Muramidase Ab-1 (RB-372-A, Thermo Fisher Scientific, UK, 1:300), Ki-67 Monoclonal Antibody (MA5-14520, Thermo Fisher, 1:150), Anti-DCLK1 antibody (ab31704, Abcam, Uk, 1:600) and Anti-Keratin 20 antibody (D9Z1Z) (ab13063, Cell Signaling Technology, 1:500). Slides were then washed in TBST and incubated with Dako envision anti-rabbit or anti-mouse labelled polymer-HRP (Dako, USA) secondary antibody for 30 minutes at room temperature, washed in TBST and incubated with 3, 3-diaminobenzidine (DAB) solution (Dako, USA) until signal was detected. Image analysis was performed using the Aperio ImageScope software v12.0.1.5027, using the specifically designed Nuclear v9.1 or Positive Pixel Count v9.1 algorithms. The algorithm was tested on three separate slides before being applied to all slides to be analysed.

### Immunofluorescence

Sections prepared as above were stained overnight at 4°C with the following antibodies: rabbit anti-Keratin 14 (Biologend, USA 905304, 1:500), Purified anti-mouse Keratin 6A (Biolegend, 905701, 1:500) and Purified anti-Loricrin Antibody (Biolegend, 905104, 1:500). The following day, sections were washed in TBST and incubated with Alexa Fluor® 594 AffiniPure Goat Anti-Rabbit IgG (H+L) (Abacus DX, Australia) with Spectral DAPI (FP1490) added (1/1000) (Perkin Elmer, USA) for one hour at room temperature. Sections were then washed in TBST and mounted using DPX/Vectashield mounting media (Vector Laboratories, Burlingame, CA). Stained slides were scanned on an Olympus IX-81 inverted fluorescence microscope slide scanner (Olympus Corporation, Japan) and imaged using the HALO software v3.1.1076.

### Intestinal epithelial cell collection

For isolation of intestinal epithelial cells, mice were euthanized and the small intestine (duodenum, jejunum and ileum) and colon removed and opened longitudinally. The tissue was washed in warm PBS and incubated in pre-warmed Buffer 1 (PBS 93%, 0.5M EDTA 3%, RNA secure 4%, Thermo Fisher Scientific, Australia), at 37°C on a shaker for 10 minutes. Samples were then vortexed, the smooth muscle removed, and the epithelial cells pelleted by centrifugation at 1400g for 5 minutes at 4°C. The pellet was washed in 1-2mL of Buffer 2 (PBS 96%, RNA secure 4%), centrifuged at 17,000 rpm for 1 minute, and pellets snap-frozen on dry ice and stored at −80°C.

### RNA isolation and quantitative real time polymerase chain reaction (q-RT-PCR)

RNA was extracted from isolated epithelial cells or tissue using the ReliaPrep RNA Tissue Miniprep System (Promega, USA) as per manufacturer’s instructions. RNA (500 ng) was converted to cDNA using the High-Capacity cDNA Reverse Transcription Kit (Applied Biosystems, USA) as per manufacturer’s instruction using random primers. Relative gene expression was determined by quantitative real time PCR (q-RT-PCR) using the ViiA 7 Real Time PCR system (Applied Biosystems, USA). The reaction mixture consisted of 0.75μL 500nM forward primer, 0.75μL 500nM reverse primer, 1μL of 1:10 diluted cDNA, 2.5μL 2x Power SYBR Green PCR Master Mix (Thermo Fisher Scientific, Australia). Samples were amplified using the following program: 95°C for 10 minutes followed by 40 cycles of 95°C for 15 seconds and 60°C for 1 minute. Primers used for q-RT-PCR are provided in **Table S1**.

### RNA-seq analysis

RNA isolated from colonic epithelial cells was prepared for next-generation sequencing using the stranded RNA library preparation kit with rRNA depletion (Illumina) and sequenced using an Illumina HiSeq 2500 with 100bp single reads. Library preparation, sequencing and initial data analysis (read mapping) was performed by an external service provider (Australian Genome Research Facility WEHI, Australia). Gene set enrichment analyses were performed using the publicly available GSEA v4.1.0 software from the Broad institute (Subramanian et al., 2005), by comparison to the MSigDB “HALLMARKS” gene set signatures (Liberzon et al., 2015). The “Signal2Noise” metric was used for gene ranking and the threshold for false discovery rate (FDR) was set at q<0.05.

### Dextran sodium sulphate treatment

Mice (n=8 per group, equal number of males and females) were given dextran sodium sulphate (DSS, 2.5% w/v, Lot#Q8378, MP Biomedicals, Canada) in autoclaved drinking water *ab libitum* for 5 days. A control group was included in which n=4 mice per genotype (equal number of males and females) were given autoclaved water without DSS for 5 days. During the experimental period, mice were weighed and monitored daily. Two hours prior to cull all mice were given an intraperitoneal injection of BrdU solution (10μL/g mouse). The disease severity score was assessed by computing the cumulative score from the following 4 criteria: (1) Weight loss: 0: Normal, 1: <5%, 2: 6-10%, 3: 11-20%, 4: >20%; (2) Faeces: 0: Normal, 1: Pasty, semi-formed, 4: Liquid, sticky or unable to defecate >5 minutes when stimulated; (3) Blood: 0: No blood, 1: Traces of blood in the stool or the rectum, 2: Free-flowing blood from the rectum or blood on fur; and (4) General appearance: 0: Normal, 1: Piloerection, hunching, 2: Lethargy and piloerection, 4: Motionless.

### Statistical analysis

Statistical analyses were performed using GraphPad Prism v8.0 software (GraphPad Software, La Jolla, CA, USA). Groups were compared using the student’s T-test with Welch’s correction or Chi-square analysis. In all cases *P*<0.05 was considered statistically significant.

## Results

### Whole body Ehf knockout mice are viable and fertile

To induce *Ehf* deletion in the whole animal *Ehf*^*Lox/+*^ mice were crossed to CMV^Cre^ mice to induce Cre-mediated recombination in all tissues, including germ cells. Resulting *Ehf^+/−^* progeny lacking *CMV^Cre^* were then inter-crossed to generate constitutive *Ehf*^*−/−*^ mice, and the colony maintained by *Ehf* heterozygous matings. Genotyping of multiple tissues confirmed systemic *Ehf* deletion (**Figure S1A**). *Ehf* mRNA expression was also examined across multiple tissues using primers specific for exon 8 encoding the floxed Ets domain. In *Ehf^+/+^* mice *Ehf* mRNA expression was highest in the salivary and preputial glands, prostate, colon and stomach, while no expression was detected in any tissues in *Ehf*^*−/−*^ mice (**Figure S1B**). Notably, the 5’ portion of the EHF transcript was expressed in *Ehf*^*−/−*^ mice at similar levels to *Ehf^+/+^* control mice raising the possibility that non-DNA binding related functions of EHF protein may be retained (**Figure S1C**).

To assess the impact of homozygous *Ehf* deletion on viability during embryogenesis, the genotype of 658 pups generated from 100 litters born to *Ehf^+/−^* x *Ehf^+/−^* breeders (average 6.6 pups per litter) was investigated. Of the 649 pups successfully genotyped, 175 were *Ehf^+/+^*, 332 were *Ehf^+/−^*, and 142 were *Ehf*^*−/−*^. The slightly lower percentage (21.9%) of *Ehf*^*−/−*^ mice born compared with the expected Mendelian percentage (25%) was not statistically significant (*P*=0.314, two-sided χ^2^ test), indicating constitutive *Ehf* deletion does not result in embryonic lethality.

### Ehf^−/−^ mice have reduced weight gain and develop multiple pathologies

To assess the impact of *Ehf* deletion on animal health, *Ehf*^*−/−*^ and *Ehf^+/+^* mice were routinely monitored for 12 months, including weekly body weight measurements for the first 27 weeks. Male *Ehf*^*−/−*^ mice weighed less than their wildtype littermates at 4 weeks, and this difference became progressively more pronounced as the mice aged (**Figure 1A**). Female *Ehf*^*−/−*^ mice weighed the same as wildtype littermates at 4 weeks, but subsequent weight gain was significantly less than wildtype littermates (**Figure 1B**). The reduced body weight of *Ehf*^*−/−*^ female mice was not due to differences in daily food intake, which was similar between genotypes (**Figure S2**).

**Figure 1.**
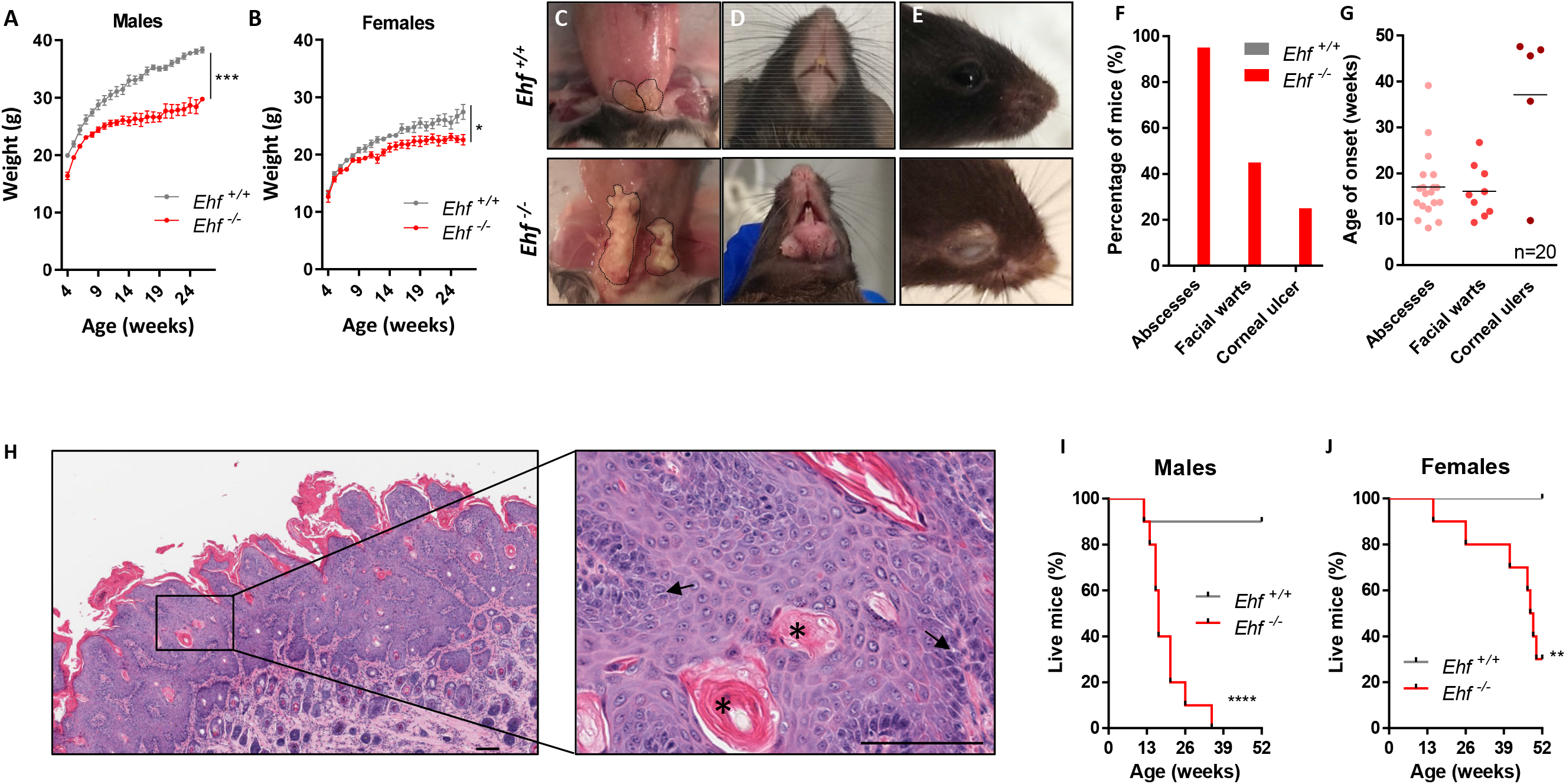
Impact of Ehf deletion on body weight and animal health. (**A, B**) Body weight of (**A**) male and (**B**) female *Ehf^+/+^* and *Ehf*^*−/−*^ mice, 4-27 weeks of age. Values shown are mean ± SEM of n=4 mice per genotype. (**C-E**) Representative images of (**C**) enlarged preputial gland abscesses, (**D**) facial papules, and (**E**) corneal ulcers in *Ehf*^*−/−*^ mice. Corresponding regions in *Ehf^+/+^* mice are shown for comparison. (**F**) Percentage of *Ehf^+/+^* and *Ehf*^*−/−*^ mice that develop preputial gland/vulval abscesses, facial lesions and corneal ulcers from analysis of n=20 mice per genotype. (**G**) Distribution of age of onset of abscesses, facial lesions and corneal ulcers from analysis of n=20 Ehf^−/−^ mice. (**H**) Haematoxylin and eosin stain of a facial papule from *Ehf*^*−/−*^ mice. Scale bar = 100 μm. Keratinous pearls are indicated by asterisks (*) and intercellular bridging by arrows. (**I, J**) Ethically-determined survival of (**I**) male and (**J**) female *Ehf^+/+^* and *Ehf*^*−/−*^ mice. Mice were censored after 52 weeks. Curves were generated from n=10 mice per genotype.

Ehf^−/−^ mice developed a series of pathologies as they aged, the most prominent being the onset of suppurative inflammation and abscesses of the preputial glands (9/10 male mice) or vulva (10/10 female mice). Consistent with this etiology, growth of the opportunistic bacteria *Staphylococcus aureus* and *Proteus mirabilis* was detected in bacterial isolates from these lesions. A high proportion of *Ehf*^*−/−*^ mice (45%) also developed papules, swelling, and ulcers around the chin and mouth which would resolve then reoccur, while 25% of mice also developed corneal ulcers. Comparatively, none of these pathologies were observed in *Ehf^+/+^* mice (**Figure 1C-F**). The median age of onset of genital abscesses and facial papules was 16 weeks, while the corneal ulcers presented later with a median age of onset of 36 weeks (**Figure 1G**). Histological analysis of the papules that developed around the chin and mouth of *Ehf*^*−/−*^ mice revealed them to be papillomas with evidence of hyperplasia, hyperkeratosis and immune cell infiltration. Histological features reminiscent of squamous cell carcinoma were also evident including keratinous pearls, enlarged nuclei and intercellular bridging (**Figure 1H**). PCR testing for polyoma virus returned a negative result suggesting a non-viral etiology.

The development of these pathologies resulted in the majority of *Ehf*^*−/−*^ mice reaching an ethical endpoint within 12 months (**Figure 1I, J**). The most common reason for euthanasia was the development of abscesses in the preputial glands and vulvae or restricted urination, in 20% (4/20) and 40% (8/20) of mice respectively. The resulting lifespan was 48 weeks for *Ehf*^*−/−*^ female mice and only 17 weeks for *Ehf*^*−/−*^ male mice, which was due mostly to the greater severity of the preputial gland abscesses compared to those which arose in the vulvae.

### Ehf deletion disrupts epidermal morphology in adult mice

Given the development of skin pathologies in *Ehf*^*−/−*^ mice, and previous findings that EHF plays a role in basal keratinocyte differentiation *in vitro* (Rubin et al., 2017), we examined the impact of Ehf deletion on normal skin histology. No difference in the histology of the chin skin was observed between *Ehf^+/+^* and *Ehf*^*−/−*^ mice at 6 weeks, with the epidermis consisting of a well-defined single cell basal layer and a thin layer of the stratum corneum in both cases (**Figure 2A**). Comparatively, at 27 weeks, while the chin skin of *Ehf^+/+^* mice was normal, (3/5) *Ehf*^*−/−*^ mice displayed areas of hyperplasia and hyperkeratosis despite not displaying macroscopic facial papules (**Figure 2A**), while a further subset developed Munro’s microabscesses (1/5) and small keratinous pearls (2/5).

**Figure 2.**
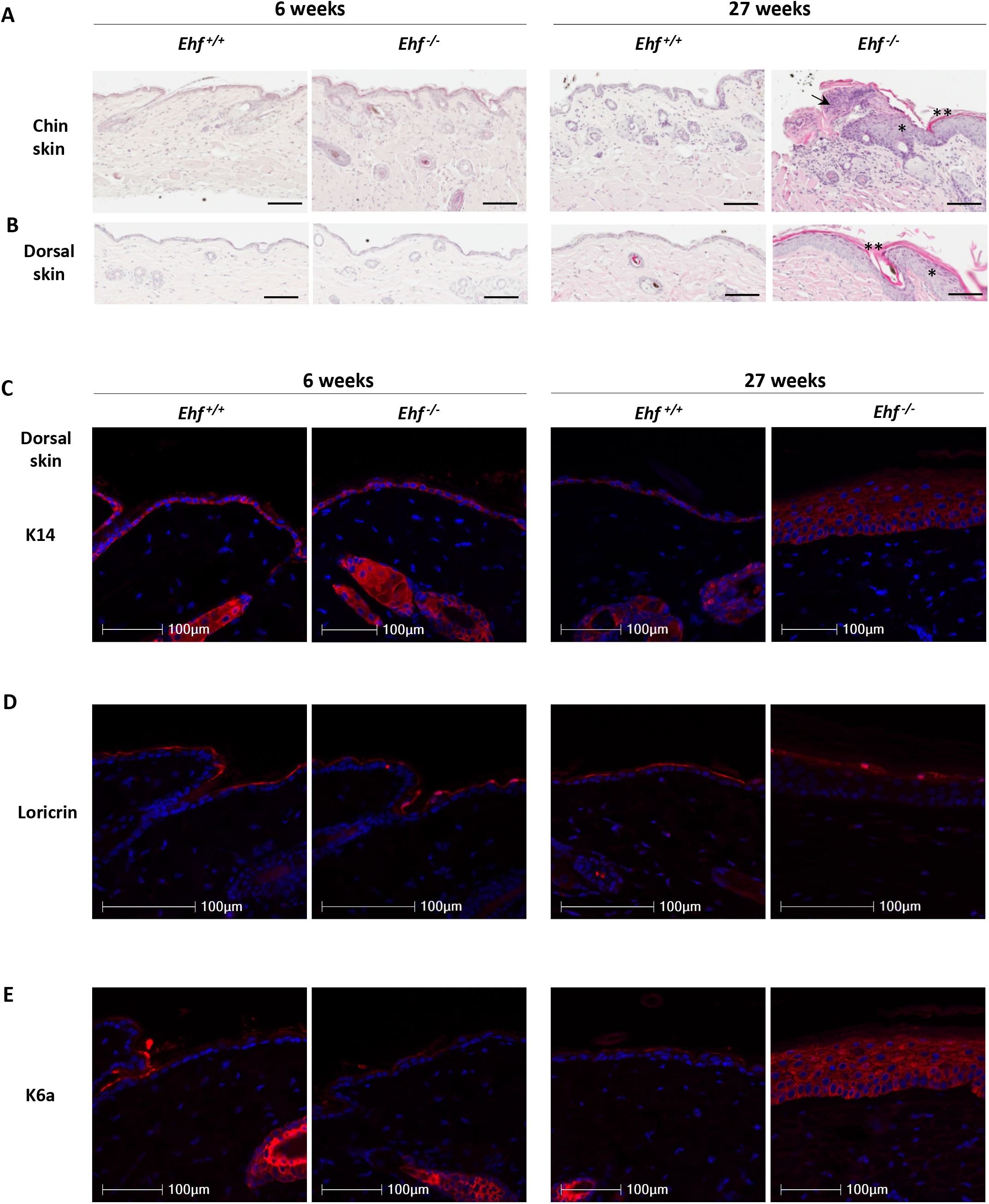
Impact of *Ehf* deletion on epidermal histology. (**A, B**) Representative haematoxylin and eosin stained sections showing the histology of the skin taken from the (**A**) chin or (**B**) dorsal region of *Ehf^+/+^* and *Ehf*^*−/−*^ littermates at 6 and 27 weeks. * Hyperplasia, ** Hyperkeratosis, Arrow: Munro’s microabscess. Scale bar = 300μm. (**C-E**) Immunofluorescence staining for (**C**) K14, (**D**) Loricrin and (**E**) K6a in the dorsal skin of *Ehf^+/+^* and *Ehf*^*−/−*^ mice at 6 and 27 weeks.

Assessment of the dorsal skin of these mice also revealed no difference in histology at 6 weeks (**Figure 2B**). At 27 weeks the dorsal skin of *Ehf^+/+^* mice was largely normal, although one mouse displayed mild hyperkeratosis and another developed Munro’s microabscesses. Comparatively, the majority of *Ehf*^*−/−*^ mice (4/5) displayed abnormal dorsal skin histology with varying degrees of hyperkeratosis and hyperplasia. Two of these mice also developed Munro’s microabscesses and wounds accompanied by immune cell infiltrates in the dermal and epidermal skin layers (**Figure 2B**).

We next stained the dorsal and chin skin for keratin 14 (K14) and Loricrin to determine whether keratinocyte differentiation was impaired. Clear staining of K14 in the basal keratinocyte layer, and of loricrin in the stratum corneum was observed in both *Ehf^+/+^* and *Ehf*^*−/−*^ mice (**Figure 2C, D**), demonstrating that *Ehf* deletion does not impact on the capacity of basal keratinocytes to differentiate into corneocytes. We also stained the dorsal and chin skin with keratin 6a (K6a), which is typically expressed in hair follicles but also during epidermal hyperproliferation as part of the wound healing response. At 6 weeks there was no K6a staining in the epidermis however, at 27 weeks, clear K6a staining was observed in areas of damaged skin in *Ehf*^*−/−*^ mice demonstrating this feature of epidermal regeneration is also not impacted by *Ehf* deletion (**Figure 2E**).

### Ehf deletion does not impact the morphology of the prostate, salivary gland, preputial gland or gastric epithelium

As EHF is highly expressed in the prostate, salivary gland, preputial gland and stomach epithelium (**Figure S1**) we undertook a gross histological assessment of these tissues in 6 weeks old *Ehf^+/+^* and *Ehf*^*−/−*^ mice. No major histological abnormalities were observed in these tissues *Ehf*^*−/−*^ mice (**Figure S3**) indicating loss of *Ehf* does not impact the gross morphology of these tissues, at this time point.

### Ehf deletion impacts intestinal cell proliferation and differentiation

As EHF is also highly expressed in the colonic epithelium, we next investigated the impact of *Ehf* deletion on proliferation and differentiation during homeostatic renewal of this tissue. While the overall architecture of the colonic epithelium remained similar between *Ehf^+/+^* and *Ehf*^*−/−*^ mice, Ki67 staining revealed a significant expansion of the proliferative colonic crypt compartment in *Ehf*^*−/−*^ mice (**Figure 3A, B**). To investigate effects on intestinal cell differentiation, expression of markers of the major intestinal epithelial cell lineages were investigated by immunohistochemistry. Staining and quantitation of the enterocyte marker KRT20 (**Figure 3C, D**), the enteroendocrine cell marker chromogranin (Figure 3E, F) and the tuft cell marker DCLK1 (**Figure 3G H**), revealed no differences between *Ehf^+/+^* and *Ehf*^*−/−*^ mice, indicating *Ehf* deletion has little impact on the differentiation and maturation of these lineages. In contrast, *Ehf* deletion caused a significant decrease in the number of goblet cells in the colon as demonstrated by reduced PAS/AB staining (**Figure 3I, J**).

**Figure 3.**
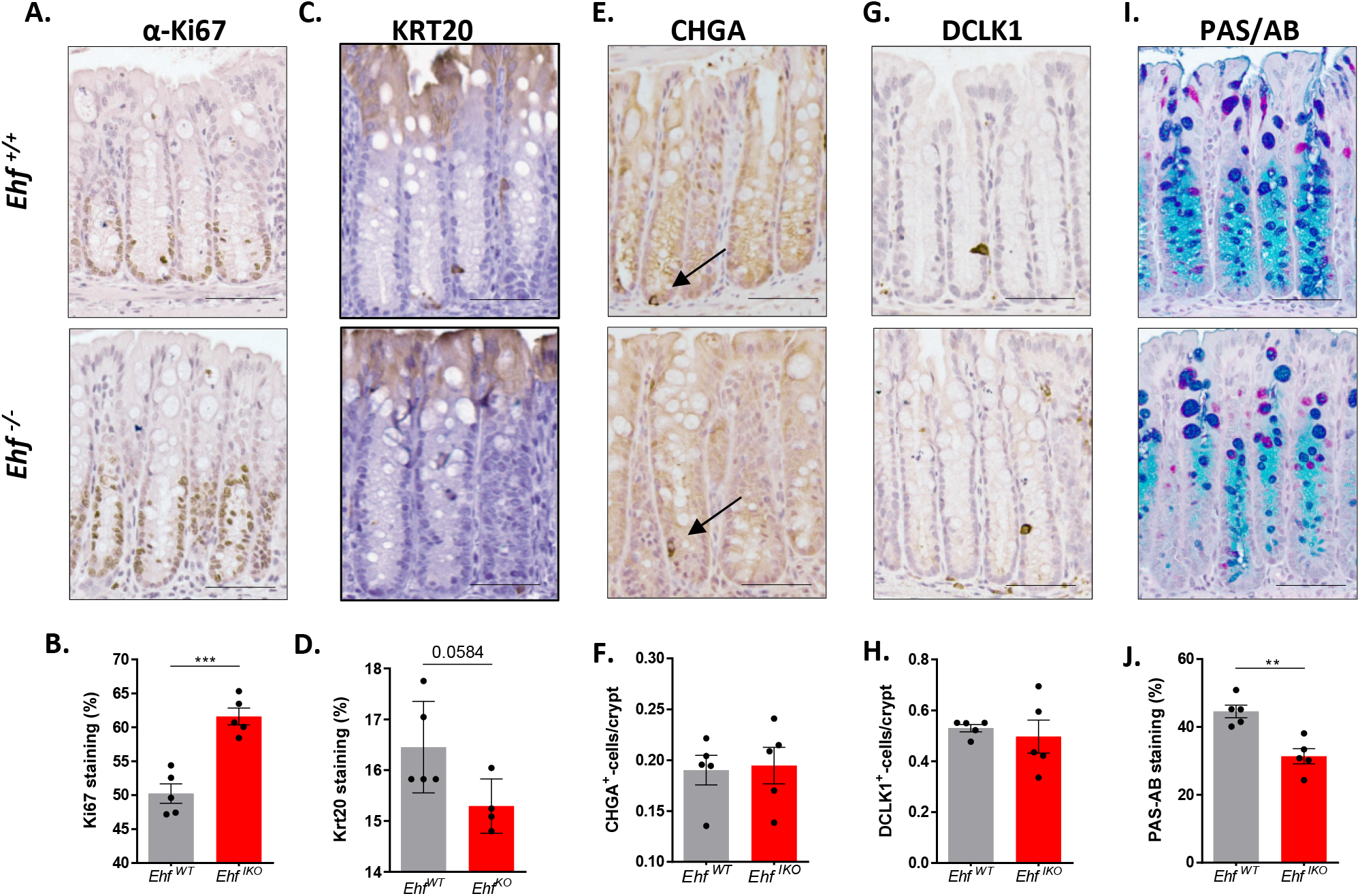
Impact of *Ehf* deletion on cell proliferation and lineage-specific differentiation in the colonic epithelium. (**A**) Staining and (**B**) quantitation of the cell proliferation marker Ki67. Staining and quantitation of the (**C, D**) enterocyte marker KRT20; (**E, F**) the neuroendocrine marker chromogranin A (CHGA); (**G, H**) the tuft cell marker DCLK1; and (**I, J**) goblet cells using periodic acid-Schiff/alcian blue (PAS/AB) staining in the colonic epithelium of 6-week-old *Ehf^+/+^* and *Ehf*^*−/−*^ mice. Scale bar = 100μm. Values shown in panels **B, D, F, H** and **J** are mean ± SEM of n=5 mice per genotype.

### Ehf deletion increases susceptibility to dextran sodium sulphate-induced colitis

Given the impact of *Ehf* deletion on colonic cell proliferation and differentiation, we next compared the response of *Ehf*^*−/−*^ and *Ehf^+/+^* mice to acute dextran sodium sulphate (DSS)-induced colitis. Weight loss was significantly more pronounced in *Ehf*^*−/−*^ compared to *Ehf^+/+^* mice over the 5-day treatment period (**Figure 4A**). Furthermore, the disease severity score was 2.5-fold higher in *Ehf*^*−/−*^ compared to *Ehf^+/+^* mice (**Figure 4B**). In addition, 25% of *Ehf*^*−/−*^ mice displayed hunching behaviour, indicative of discomfort and/or pain at the end of the experimental period while all *Ehf^+/+^* mice displayed normal behaviour. Consistent with these changes, the length of the colon was significantly shortened in DSS-treated *Ehf*^*−/−*^ mice (**Figure 4C,D**), and histological analysis of the colon revealed that *Ehf* deletion increased the extent and depth of immune cell infiltration, increased the area of crypt loss and ulceration, and led to more widespread oedema (**Figure 4E-M**).

**Figure 4.**
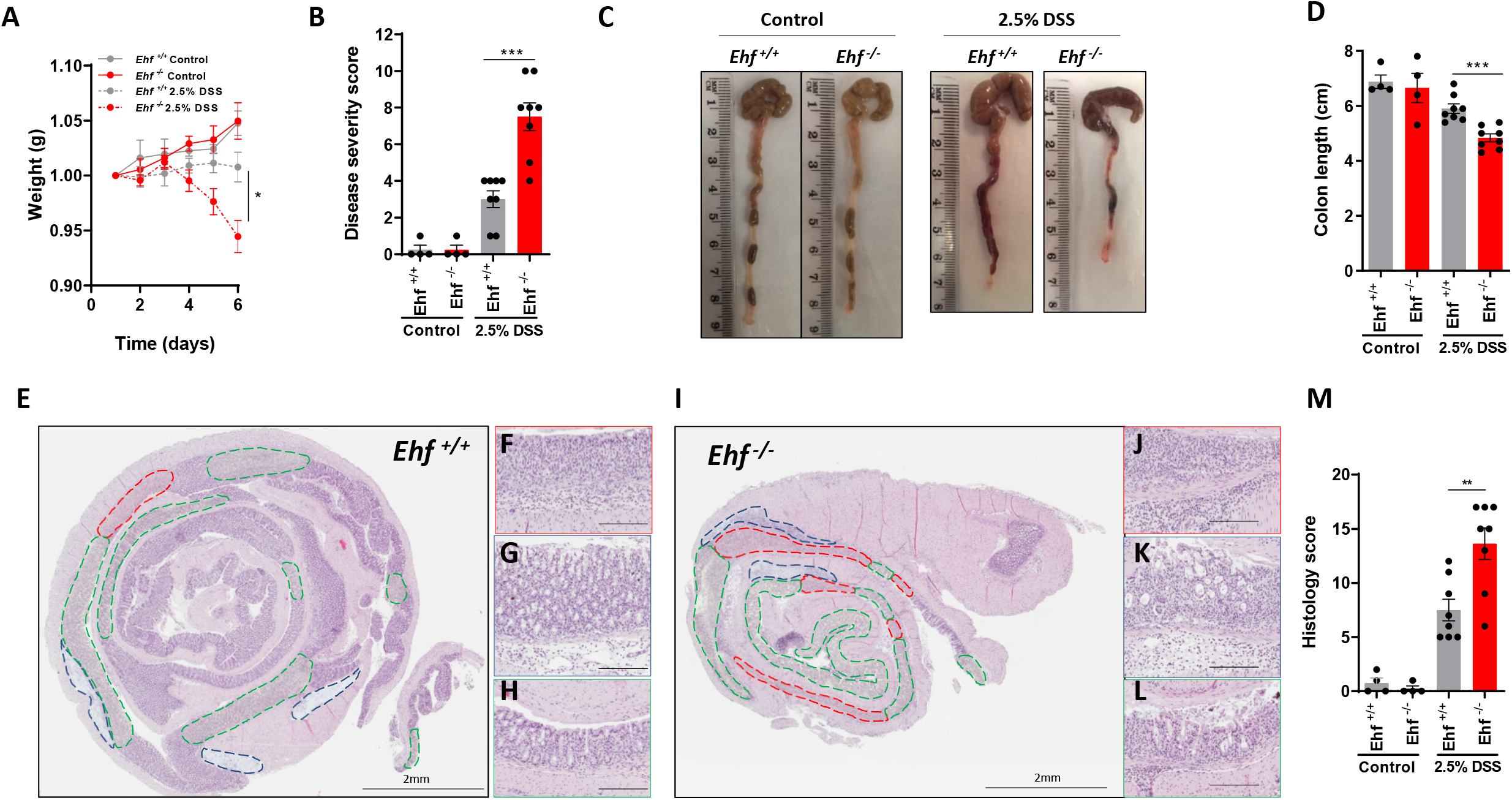
Sensitivity of *Ehf*^*−/−*^ mice to DSS-induced colitis. (**A**) Change in body weight of *Ehf^+/+^* and *Ehf*^*−/−*^ mice with or without treatment with 2.5% DSS. (B) Summary of the disease severity score of *Ehf^+/+^* and *Ehf*^*−/−*^ mice treated with or without 2.5% DSS. (**C**) Representative images of the colon lengths of *Ehf^+/+^* and *Ehf*^*−/−*^ mice treated with or without 2.5% DSS and (D) corresponding quantitation of these data. (**E, F**) Representative haematoxylin and eosin stained sections of the entire colon from *Ehf^+/+^* and *Ehf*^*−/−*^ mice treated with 2.5% DSS. Regions of ulceration are highlighted by a red border, areas of oedema with a blue border, and areas of epithelial erosion and/or crypt loss with a green border. (**F-H, J-L**) High power fields of regions of (**F, J**) ulceration, (**G, KL**) oedema, and (**H, L**) epithelial erosion and/or crypt loss. Scale bar = 200μm. (**M**) Summary of the histology scores from *Ehf^+/+^* and *Ehf*^*−/−*^ mice treated with or without 2.5% DSS. Values shown in **A, B, D** and **M** are mean ± SEM of n=4 and n=8 mice per genotype in the control and DSS-treated groups respectively.

### Intestinal-specific Ehf deletion increases sensitivity to DSS-induced colitis

To confirm that these effects were due to alterations in the intestinal epithelium, we generated intestinal-specific *Ehf* knockout mice (*Ehf*^*IKO*^) by crossing *Ehf*^*Lox/Lox*^ mice to *Villin^CreERT2^* mice, and treatment of the *Ehf*^*Lox/Lox*^; *Villin^CreERT2^* offspring with tamoxifen. Genotyping of small intestinal epithelial cells isolated from 6-week-old *Ehf*^*IKO*^ or sunflower oil treated controls (*Ehf*^*WT*^) confirmed selective deletion of Exon 8 of *Ehf* in the tamoxifen treated group (**Figure S4A**). As expected, *Ehf* mRNA levels were significantly down-regulated in small intestinal and colonic epithelial cells in *Ehf*^*IKO*^ but not *Ehf*^*WT*^ mice (**Figure S4B**). The absence of *Ehf* mRNA was also examined in 6-month-old *Ehf*^*IKO*^ mice which confirmed sustained deletion of *Ehf*, and that no compensatory repopulation of the intestinal epithelium by *Ehf* WT cells occurs over time (**Figure S4C**).

We next assessed the susceptibility of *Ehf*^*IKO*^ mice to DSS-induced colitis. As observed in whole body *Ehf*^*−/−*^ mice, the onset of colitis was markedly more severe in *Ehf^IKO^* compared to *Ehf*^*WT*^ mice, assessed by both the disease severity (**Figure S4D**) and histology scores (**Figure S4E-M**), confirming EHF-mediated disruption of colonic homeostasis is epithelial driven.

### Intestinal-specific Ehf deletion induces extensive transcriptional reprogramming in the colonic epithelium

To further elucidate the role of EHF in the colonic epithelium, we performed RNA-seq analysis of colonic epithelial cells isolated from 6-week-old *Ehf*^*WT*^ and *Ehf*^*IKO*^ mice. To account for possible transcriptional changes induced by tamoxifen, *Villin^CreERT2^* mice treated with tamoxifen (*Villin*^*CreERT2-TMX*^) were also analysed as a further control. Principal components analysis of the RNA-seq data resulted in clear separation of the *Ehf*^*IKO*^ samples from the two control groups (**Figure S5**), indicating *Ehf* deletion induces extensive transcriptional reprogramming in the colonic epithelium. Analysis of the gene expression changes identified 146 differentially expressed genes between the *Ehf*^*IKO*^ mice and both control strains, of which 75 were up-regulated and 71 were down-regulated in the *Ehf*^*IKO*^ mice (**Figure 5A**). Of the top differentially expressed genes, increased expression of REG4, ID4 and PLA2G21, and reduced expression of LPO in *Ehf*^*IKO*^ mice was confirmed by q-RT-PCR demonstrating the reproducibility of the RNA-seq data (**Figure 5B**).

**Figure 5.**
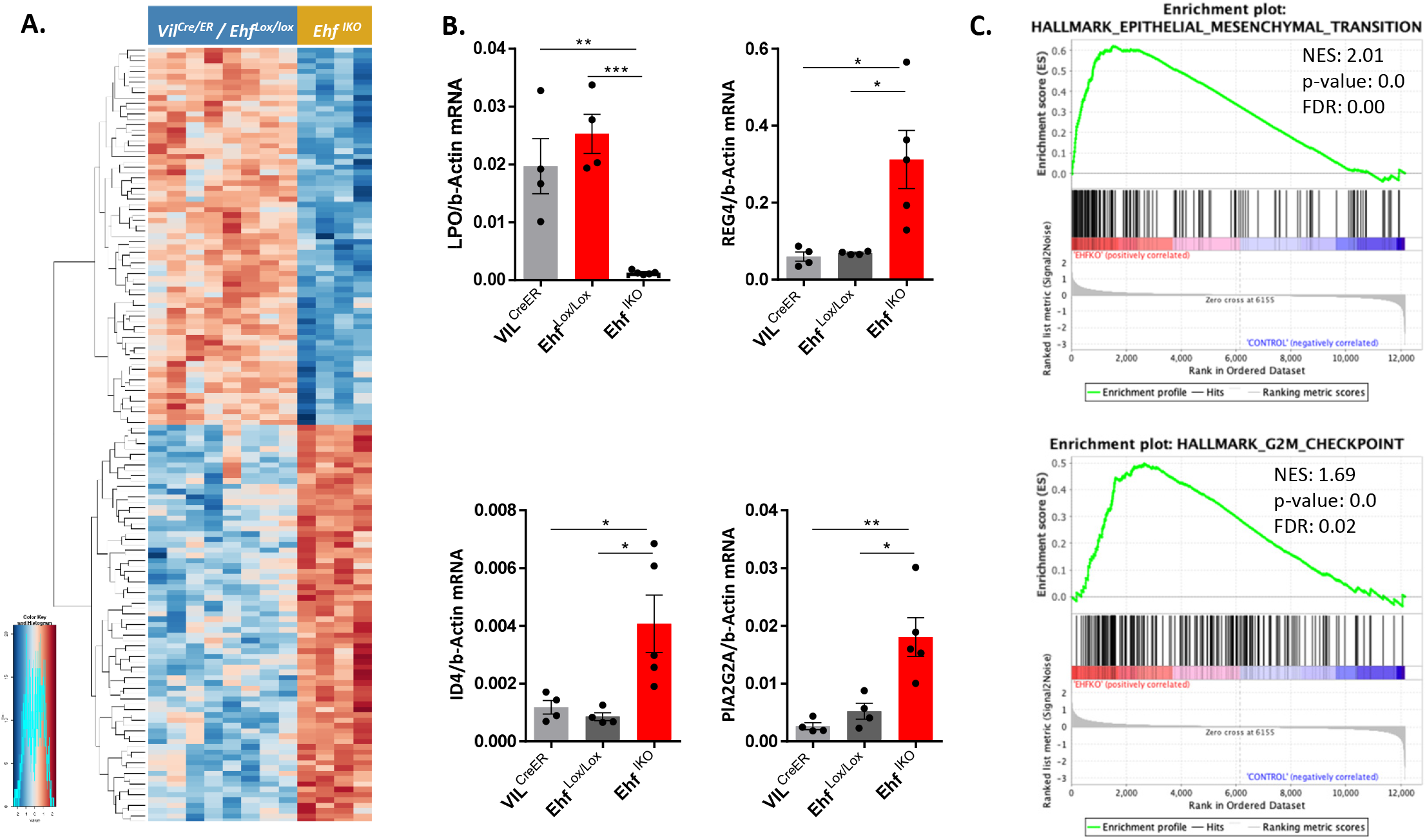
*Ehf* deletion induces extensive transcriptional reprogramming of the colonic epithelium. (**A**) Heatmap of genes differentially expressed between *Ehf^IKO^* and both Ehf^Lox/Llox^ and *Villin^CreERT2-TMX^* mice. (**B**) q-PCR validation of selected differentially expressed genes. Values shown are mean ± SEM of n=4, n=4 and n=5 for *Villin^CreERT2-TMX^*, *Ehf^Lox/Llox^* and *Ehf^IKO^* mice respectively. (**C**) Gene set enrichment plots showing significant enrichment of the epithelial to mesenchymal transition (146 genes) and G2M checkpoint (192 genes) hallmarks in *Ehf^IKO^* mice. NES = normalized enrichment score, FDR = False discovery rate q-value.

Consistent with the increase in colonic cell proliferation induced by *Ehf* deletion (**Figure 3A**), gene set enrichment analysis (GSEA) identified an enrichment of the G2M checkpoint genes, which included increased expression of E2F3, CENPF and RAD21 in *Ehf*^*IKO*^ mice (**Figure 5C, Table S2**). The GSEA also identified an enrichment of the epithelial to mesenchymal transition (*EMT*) gene signature, which included increased expression of *VCAM1, ITGB3, FAP, FBN2, TFPI2, ADAM12, ELN, WNT5A, SPP1, BGN, SLIT3, SERPINH1, FBLN2, PLAUR, CAPG, TNC, TIMP3, LAMC1, PMP22* and *ITGB5* and several collagens (*COL1A1, COL1A2, COL3A1, COL4A1, COL4A2, COL5A1, COL5A2, COL6A2, COL6A3*) in *Ehf*^*IKO*^ mice (**Figure 5C, Table S3**).

### EHF protects against Apc-initiated colonic adenoma development

Finally, to extend these findings, we examined the impact of *Ehf* deletion on tumour development specifically in the colon. To achieve this, *Ehf*^Lox/Lox^ mice were crossed to Cdx2^*CreERT2*^ deleter mice (*Ehf*^*CKO*^), and in turn to *Apc*^*Lox/+*^ mice, to induce compound homozygous deletion of *Ehf* and heterozygous deletion of Apc (*Apc*^*CKO/+*^) in the epithelium of the colon and caecum. *Ehf*^*CKO*^; *Apc*^*CKO/+*^ and *Ehf*^*WT*^; *Apc*^*CKO/+*^ controls were subsequently monitored for colon tumour development over 52 weeks. While there was no difference in ethically determined survival, *Ehf*^*CKO*^ *Apc*^*CKO/+*^ male mice had significantly lower body weight compared to *Ehf*^*WT*^; *Apc*^*CKO/+*^ mice, although this difference was not observed in female mice (**Figure S6A-C**). At the end of the 52-week experimental period, mice were culled and tumours quantified. Notably, more *Ehf*^*CKO*^; *Apc*^*CKO/+*^ mice (6/11) developed adenomas compared with *Ehf*^*WT*^; *Apc*^*CKO/+*^ mice (4/12) and the number of colonic adenomas per mouse was higher in the *Ehf*^*CKO*^; *Apc*^*CKO/+*^ mice, however this difference was not statistically significant (**Figure 6A**). Comparatively, colonic tumour burden was significantly higher in *Ehf*^*CKO*^ *Apc*^*CKO/+*^ mice compared with *Ehf*^*WT*^; *Apc*^*CKO/+*^ mice, demonstrating that EHF suppresses the growth of Apc-initiated colonic adenomas (**Figure 6B, C**). We also observed that while adenomas arose predominantly in the caecum in *Ehf*^*WT*^; *Apc*^*CKO/+*^ mice, a higher proportion of tumours in the proximal, middle and distal colon were observed in Ehf^CKO^ *Apc*^*CKO/+*^ mice, demonstrating EHF preferentially suppresses tumour formation in these regions (**Figure 6D**).

**Figure 6.**
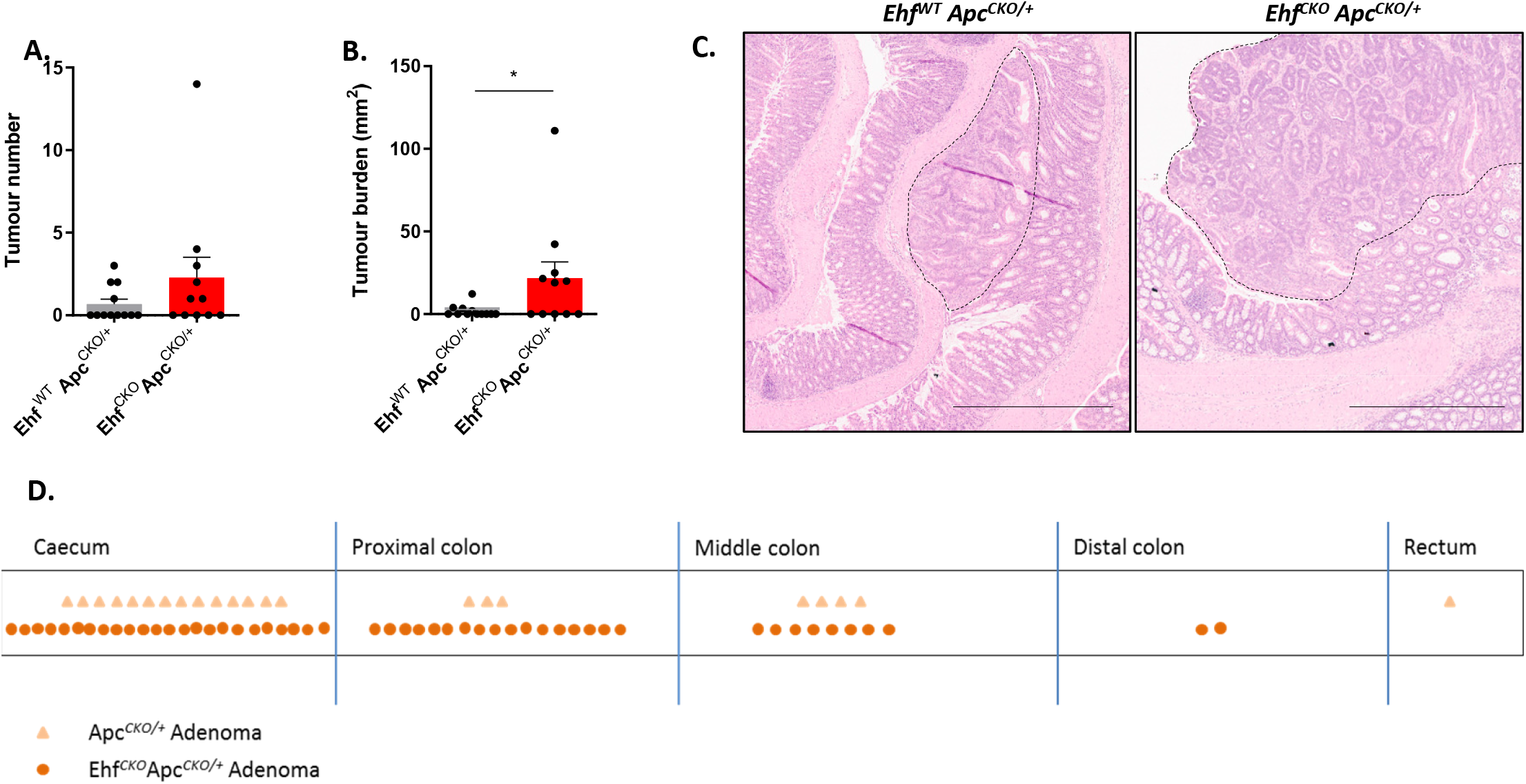
EHF protects against *Apc*-initiated tumour formation in the colon. (**A, B**) Effect of Ehf deletion on (**A**) tumour number and (**B**) tumour burden. Values shown are mean ± SEM of n=12 and n=11 for *Ehf^WT^*; *Apc*^*CKO/+*^ and *Ehf^CKO^*;*Apc*^*CKO/+*^ mice respectively. (**C**) Representative H&E stained images of adenoma development in the colon of *Ehf^WT^*; *Apc*^*CKO/+*^ and *Ehf*^*CKO*;^*Apc*^*CKO/+*^ mice. (**D**) Distribution of adenomas in the ceacum and colon of *Ehf^WT^*; *Apc*^*CKO/+*^ and *Ehf^CKO^*;*Apc*^*CKO/+*^ mice.

## Discussion

By generating a novel whole body knockout mouse lacking the Ets DNA binding domain of EHF, we demonstrate an essential role for the EHF transcription factor in the maintenance of epidermal and intestinal epithelial cell homeostasis.

While *Ehf*^*−/−*^ mice are born at a slightly lower than expected Mendelian ratio, this difference was not significant, establishing that EHF is not essential for embryonic development. This contrasts with the whole animal knockout phenotype of the related ESE family member *Elf5* which results in lethality by embryonic day 7.5 (Donnison et al., 2005; Zhou et al., 2005), and *Elf3* which results in embryonic lethality in approximately 30% of mice (Ng et al., 2002). Comparatively, the viability of *Ehf*^*−/−*^ mice at birth is similar to that of whole body Spdef knockout mice which are born at expected Mendelian ratios (Gregorieff et al., 2009; Horst et al., 2010).

In our model we specifically deleted the major known function of EHF, it’s DNA binding and transcriptional activity, by targeted deletion of the Ets DNA binding domain. It is possible to conclude therefore that the effects observed on epidermal and colonic epithelial homeostasis are a consequence of loss of EHF transcriptional activity. It is also possible that some DNA binding-independent functions of EHF may be retained in this model, although this is yet to be confirmed. For instance, the pointed domain of Ets proteins mediates protein-protein interactions, and while it was initially shown that isolated pointed domains from Ets transcription factors did not retain functionality (Mackereth et al., 2004), a recent study of SPDEF demonstrated functionality of the pointed domain independent of the Ets domain (Lo et al., 2017). Notably, when the Ets domain was deleted, the pointed domain of SPDEF was able to bind and sequester β-catenin in the cytoplasm more effectively. Comparatively, deletion of the Ets DNA binding domain of *Elf5* resulted in complete loss of ELF5 protein (Choi et al., 2009). Despite extensive effort, we were unable to identify an antibody which could reliably detect mouse EHF protein by western blot or IHC. We therefore cannot rule out the possibility that some functionality of EHF may be maintained in *Ehf*^*−/−*^ mice. Nevertheless, our current findings establish that deletion of the DNA binding domain of EHF is sufficient to induce a series of phenotypes in multiple tissues.

Most notable among these was the reduced body weight of both male and female *Ehf*^*−/−*^ mice compared to *Ehf^+/+^* littermates, and the development of abscesses in the preputial glands/vulvae, papillomas in the chin/facial skin, and corneal ulcers in *Ehf*^*−/−*^ mice. Preputial glands in mice are susceptible to bacterial infections (Bertrand et al., 2016), and once infected can cause pain, self-mutilation, restricted urination, penile prolapse and fistulation, and was the most common reason for *Ehf*^*−/−*^ male mice having to be euthanised within one year.

Examination of the epidermal layer of *Ehf*^*−/−*^ mice revealed minimal differences in 6 week old mice, and normal staining of loricrin was observed in the stratum corneum, demonstrating that *Ehf* deletion does not impact on the capacity of basal keratinocytes to differentiate into corneocytes. Nevertheless, hyperplasia, hyperkeratosis and microabscesses were observed in over 60% of mice at 27 weeks of age. The later onset of these lesions and the observation that they mostly occur in areas of high exposure to damage-inducing behaviours such as grooming and fighting, suggests a potential role for *EHF* in repair of the skin epithelium. Such a role is consistent with a recent study involving genome-wide profiling analysis of enhancer regions in human keratinocytes which identified the Ets motif as the most enriched transcription factor binding site in these cells, and revealed that EHF regulates ~400 genes in these cells including several associated with keratinocyte differentiation (Rubin et al., 2017).

Interestingly, a subset of *Ehf*^*−/−*^ mice developed corneal ulcers. Notably, a recent study identified *Ehf* as one of the few genes significantly up-regulated during ageing in the mouse corneal epithelium (Stephens et al., 2013), and ChIP analysis demonstrated EHF binding to *Tcf4*, a transcription factor required for corneal stem cell maintenance (Lu et al., 2012). EHF knockdown in corneal epithelial cells also resulted in decreased expression of SPRR which is required for keratin cross-linking and maintenance of the barrier function of the skin (Stephens et al., 2013). These findings suggest that the corneal defects observed in *Ehf*^*−/−*^ mice may be due to defective corneal differentiation during ageing.

We also observed that *Ehf*^*−/−*^ mice were more sensitive to DSS-induced colitis, which was phenocopied in gut epithelial-specific *Ehf* knockout mice, confirming this effect was epithelial cell dependent. Interestingly, we noted a reduced number of mucin-producing goblet cells in *Ehf^IKO^* mice. Given the important role of mucin in protecting the colonic epithelium from damage (Velcich et al., 2002), it is possible that the reduced number of goblet cells may underpin the susceptibility of *Ehf^IKO^* mice to DSS-induced colitis. *Ehf* deletion also significantly decreased expression of *Spdef* and *Creb3l1* which are important determinant of the goblet cell lineage in the intestinal epithelium (Gregorieff et al., 2009; Noah et al., 2010), providing a possible mechanistic explanation for the reduction in goblet cell differentiation in these mice.

Importantly, Zhu *et al* recently reported that EHF is a downstream target of the lncRNA lncGata6 (Zhu et al., 2018), and generated a mouse with whole-body deletion of *Ehf* using CRISPR-Cas9-mediated genome editing. They reported that EHF was required for the maintenance of small intestinal stem cells by driving expression of *Lgr4* and *Lgr5*, and that *Ehf* deletion significantly decreased Olfm4-positive stem cells and reduced the number of all secretory lineage cell types in the small intestine (Paneth, tuft, enteroendocrine and goblet cells). While the focus of our study was the colonic epithelium where EHF is most highly expressed, examination of Olfm4-positive stem cells in the small intestine revealed no difference between *Ehf^+/+^* and *Ehf*^*−/−*^ mice (data not shown). One explanation for this difference may be that the *Ehf* knockout mice generated by Zhu *et al* completely lacked any functional part of the Ehf gene, as transcription was prematurely terminated in exon 2, whereas only the DNA binding domain was deleted in our model.

Notably, we observed a significant increase in cell proliferation in *Ehf*^*−/−*^ mice evidenced by increased Ki67 staining. Consistent with this finding, gene-set enrichment analyses revealed a significant enrichment of G2/M checkpoint genes in *Ehf^IKO^* mice. Whether EHF impacts cell proliferation by directly regulating expression of these genes, or indirectly by altering key colonic epithelial signalling pathways, or other mechanisms, remains to be determined. The GSEA analysis also identified a significant enrichment of genes involved in epithelial to mesenchymal transition in *Ehf^IKO^* mice. Interestingly, examination of the corresponding protein expression profile of these genes in the normal human colon using the human protein atlas database (Uhlen et al., 2015) revealed that several of these genes (*ITGB3, FBN2, TFPI2, ELN, BGN, FBLN2, CAPG, TNC, LAMC1, COL3A1, COL4A1, COL4A2 and COL6A3*) are not normally expressed in glandular colonic epithelial cells. Therefore, although these analyses were performed on isolated epithelial cells, further investigation is needed to determine whether these changes are occurring in the epithelial or stromal compartment, or both.

Consistent with the increase in cell proliferation and EMT following *Ehf* deletion, we identified a potential tumour suppressive role for EHF in the colon, as loss of *Ehf* increased Apc-initiated colonic adenoma formation and burden. The mechanisms driving this effect remain to be determined, but may be related to the increased rate of colonic cell proliferation in these mice, or epithelial to mesenchymal transition, which transcriptional profiling revealed to be altered in *Ehf^IKO^* mice. Additionally, the loss of goblet cells may also contribute to the increased tumour incidence as previously reported for *Muc2*^−/−^ mice (Velcich et al., 2002).

In summary, our findings represent the first systematic study of the role of the EHF transcription factor in development and tissue homeostasis in vivo. We reveal an essential role for EHF in the maintenance of epidermal and colonic homeostasis, and furthermore, a novel role in suppressing colonic tumour development.

## Acknowledgements

This project was supported by NHMRC project grant (GNT1107836), NHMRC Fellowships to JMM (GNT1046092) and ME (GTN1079257/GTN1173814), and the Operational Infrastructure Support Program, Victorian Government, Australia.

## Author contributions

C.M.R, R.N, I.Y.L, L.J.J, D.S.W, C.D, F.T, H.A, M.C, K.S, M.B, D.M, O.M.S, M.E, A.S.D, and J.M.M: Conducted, analysed and interpreted experiments.

C.M.R., R.N., M.E., A.S. D., J.M.M: Conceived, designed experiments and/or supervised parts of the study.

M.F.E and F.K: Contributed to the generation of essential reagents.

C.M.R., J.M.M: Wrote the manuscript.

F.K, C.D, M.C, M.F.E, O.M.S, A.S.D: Edited the manuscript.

O. M.S, A.S.D: Contributed to the generation of funding for the study.

